# Myosin essential light chain 1sa decelerates actin and thin filament gliding on β-myosin molecules

**DOI:** 10.1101/2022.03.08.483456

**Authors:** Jennifer Osten, Maral Mohebbi, Petra Uta, Faramarz Matinmehr, Tianbang Wang, Theresia Kraft, Mamta Amrute-Nayak, Tim Scholz

## Abstract

The β-myosin heavy chain expressed in ventricular myocardium and the myosin heavy chain (MyHC) in slow-twitch skeletal *soleus* muscle type-I fibers are both encoded by *MYH7*. Thus, these myosin molecules are deemed equivalent. However, some reports suggested variations in the light chain composition between *soleus* and ventricular myosin, which could influence functional parameters such as maximum velocity of shortening. To test for functional differences of the actin gliding velocity on immobilized myosin molecules we made use of the *in vitro* motility assay.

We found that ventricular myosin moved actin filaments with approx. 0.9 μm/s significantly faster than *soleus* myosin (0.3 μm/s). Unregulated actin filaments are not the native interaction partner of myosin and are believed to slow down movement. Yet, using native thin filaments purified from *soleus* or ventricular tissue, the gliding velocity of *soleus* and ventricular myosin remained significantly different. When comparing the light chain composition of ventricular and *soleus* β-myosin a difference became evident. *Soleus* myosin contains not only the “ventricular” essential light chain (ELC) MLC1sb/v, but also an additional longer and more positively charged MLC1sa. Moreover, we revealed that on a single muscle fiber level, a higher relative content of MLC1sa was associated with significantly slower actin gliding.

We conclude that the ELC MLC1sa decelerates gliding velocity presumably by a decreased dissociation rate from actin associated with a higher actin affinity compared to MLC1sb/v. Such ELC/actin interactions might also be relevant *in vivo* as differences between *soleus* and ventricular myosin persisted when native thin filaments were used.

**Summary:** Compared to the “ventricular” essential myosin light chain MLC1sb/v, the longer and more positively charged MLC1sa present in slow-twitch *soleus* muscle fibers decelerates actin filament gliding on β-myosin molecules presumably by a decreased dissociation rate from actin filaments.

## Introduction

Myriad essential cellular processes including cytokinesis, cargo transport, cell migration, and muscle contraction are driven by motor proteins such as myosins of different classes. The myosin holoenzyme consists of myosin heavy chains (MyHC) and light chains. Myosin heavy chains are composed of an N-terminal catalytic head or motor domain, an adjacent neck domain and a C-terminal tail domain of variable design and function. The catalytic motor domains produce movement and force by cyclic interactions with actin filaments powered by the chemical energy derived from ATP hydrolysis. Members of myosin class II dimerize by their coiled-coil tail domains, thus forming hexameric, two-headed protein complexes composed of two heavy chains and two pairs of myosin light chains (MLC), one essential light chain (ELC) and one regulatory light chain (RLC) associated with each MyHC.

Myosin light chains are calmodulin family members, which bind non-covalently to so-called IQ motifs located in the MyHC neck domain and are needed for the structural integrity of the myosin protein complex (Lowey *et al*., 1993b; VanBuren *et al*., 1994). MLCs can have regulatory functions on the motor activity of the myosin holoenzyme by modulating functional properties such as maximal velocity of shortening of muscle fibers or even processive behaviour (Amrute-Nayak *et al*., 2019; Greaser *et al*., 1988; Reggiani *et al*.,1997). Reports on hypertrophic cardiomyopathy associated MLC mutations also suggest functional relevance of the associated MLCs (Guhathakurta *et al*., 2017; Hernandez *et al*.,2007; Huang and Szczesna-Cordary, 2015). In class-II myosins, ELC binds the IQ motif next to the motor domain, while the RLC binds to the nearby second IQ motif (Rayment *et al*., 1993). Muscle tissue ELCs consist of 150–208 amino acids and have molecular weights ranging from approximately 17-23 kDa. They are of various length and can be grouped into shorter (also known as A2 type) ELCs such as MLC3f expressed in fast muscle fibers and longer (also known as A1 type) isoforms expressed in slow and fast skeletal muscle fibers and cardiomyocytes (Lowey and Risby, 1971).

The finding that myopathies such as hypertrophic cardiomyopathy (HCM) are often caused by point mutations in cardiac sarcomeric proteins has generated an increasing interest for these molecules (Rayment *et al*., 1995). In HCM, mainly the ventricular motor protein β-myosin heavy chain (β-MyHC) or the cardiac myosin-binding protein C are affected. Within β-MyHC, most mutations are located in the motor domain, which is responsible for cyclic interactions with actin filaments and generation of motion and force powered by ATP hydrolysis. Thus, mutations within the myosin motor can result in functional alterations of the actomyosin complex and might lead to organ malfunction. β-MyHC in ventricular myocardium is the product of the same gene (*MYH7*) on chromosome 14q1 (Epstein *et al*.,1992; Geisterfer-Lowrance *et al*., 1990), which is also expressed in slow-twitch type-I muscle fibers found in e.g. *Musculus soleus* (Lompré *et al*., 1984; Schiaffino and Reggiani, 1996). Normal and mutant β-MyHC are incorporated into the sarcomeres of both cardiac and slow-twitch skeletal muscle (Becker *et al*., 2007; Cuda *et al*., 1993; Nier *et al*., 1999; Tripathi *et al*., 2011; Yu *et al*., 1993), the latter one being more easily accessible and less subject to the adaptive responses seen in myocardium. Consequently, single *soleus* muscle fibers or myosin extracts from HCM patients have often been used to study functional effects of HCM-related β-myosin mutations compared to wild-type myosin from healthy individuals (Cuda *et al*., 1997; Kirschner *et al*., 2005; Köhler *et al*., 2002; Seebohm *et al*., 2009; Thedinga *et al*., 1999). However, besides variations of MyHC isoforms, some reports suggested variations in the myosin light chain composition between individual *soleus* muscle fibers, which could influence functional parameters like maximum velocity of fiber shortening (Greaser *et al*., 1988; Reiser *et al*., 1985; Sweeney *et al*., 1988). Additionally, the isoform composition of the myosin mechanoenzyme regarding MyHCs and MLCs can vary during development and can be shifted by various stimuli such as thyroid hormones (Biral *et al*., 1999; Sartore *et al*., 1981). This suggests that even when myosin molecules share the same heavy chain, their functional properties might differ due to differences in other components of the myosin complex such as MLCs.

Unlike shorter A2 type ELCs, A1 ELC isoforms contain an 40-45 amino acid long, positively charged N-terminal extension, which can transiently interact with acidic residues on F-actin, thereby bridging the actin filament and the myosin-II motor domain and likely modulating myosin motor mechanoenzymatic function (Andreev *et al*., 1999; Milligan *et al*.,1990; Sutoh, 1982; Sweeney, 1995; Timson *et al*., 1998). A1 ELC isoforms MLC1v and MLC1sb found in ventricular and slow-twitch skeletal muscle tissue have been shown to be identical and encoded by the same gene in mouse (Barton and Buckingham, 1985) and human (Fodor *et al*., 1989). In some slow-twitch muscle fibers, however, a second MLC1s isoform has been detected in addition to MLC1sb (Pinter *et al*., 1981; Sarkar *et al*., 1971; Weeds, 1976). Like MLC1v or MLC1sb, this isoform called MLC1sa belongs to the A1 type ELCs. MLC1sa contains a positively charged N-terminal extension, which is slightly longer than observed in MLC1sb. The presence of different slow-type ELC isoforms in slow-twitch skeletal muscle is heterogeneous across species and can vary between slow muscles in the same species or even fibers from the same muscle (Bicer and Reiser, 2004; Biral *et al*., 1982; Carraro *et al*., 1981; Staron and Pette, 1987), and is therefore not generally considered.

As several studies reported modulating effects of A1 ELC isoforms on myosin motor mechanoenzymatic function, the presence of an additional A1 ELC such as MLC1sa with a longer N-terminal extension might cause changes in functional properties such as maximal velocity of shortening. Indeed, in pig diaphragm slow muscle fibers maximal velocity of shortening was reported to be inversely related to the relative level of MLC1sa (Reiser and Bicer, 2006). However, in the studied muscle fibers functional effects upon changes in the relative level of MLC1sa were accompanied by changes in the expression of a fast A2 ELC isoform MLC1f and of a slow-type troponin-T isoform. The aims of the present study were, therefore, (i) to test whether differences between ventricular and slow *soleus* myosin regarding the speed of actomyosin chemomechanical interaction persist also in a reduced system such as the actin gliding assay. Assuming that there are differences regarding the protein isoform composition of myosin holoenzymes in ventricular and slow-twitch *soleus* muscle tissue, we aim to test (ii) whether possible differences in functional parameters of myosin holoenzymes can be attributed to the presence of the long ELC isoform MLC1sa expressed in slow-twitch muscle fibers.

We found that *soleus* myosin moved actin filaments significantly slower than ventricular myosin. Slower actin gliding was associated with the presence of MLC1sa in *soleus* myosin, and on a single *soleus* muscle fiber level, a higher relative content of MLC1sa was related with significantly slower actin gliding. To exclude artificial effects of unregulated actin filaments, we also confirmed differences in actin filament gliding velocity between myosin holoenzymes with or without MLC1sa using native thin filaments extracted from ventricular and *soleus* tissue.

## Materials and methods

### Animals

Female New Zealand bastard white rabbits, aged between 4 to 7 months were euthanized following the regulations from the German Animal Welfare Act. The animals used for the collection of *soleus* and ventricular muscle tissue were registered under §4/2004/462 or 504, and §4/10 Kraft, in accordance with the regulations existing at that time. For atrial samples New Zealand bastard white rabbits with the authorization numbers I2A291 F17478+F17313 (approved to Dr. Jan Faix, Hannover Medical School) were used.

### Protein purification

#### Myosin

Full-length myosin was purified from native *Musculus soleus* or ventricular tissue of New Zealand bastard white rabbits. First, the tissue was disintegrated using a liquid nitrogen cooled pestle and mortar, followed by myosin extraction for 20 min in extraction buffer (0.3 M KCl, 0.15 M K_2_HPO_4_, 0.01 M Na_4_P2O_7_, 1 mM MgCl_2_ and 2 mM DTT, pH 6.8). After centrifugation at 65000 rpm (Beckman Coulter, TLA 120.2) for 1 hour, the supernatant was diluted with pure water containing 2 mM DTT to precipitate myosin. After incubation for 40 minutes on ice and subsequent centrifugation for 30 min at 27000 rpm (Beckman coulter, Ti70) the pellet was resuspended in myosin buffer (0.3 mM KCl, 25 mM HEPES, 4 mM MgCl_2_, 1 mM EGTA, and 2 mM DTT). Protein concentration was determined using Bradford protein assay (Bio-Rad Laboratories, Inc., #5000006). Myosin was snap frozen over liquid nitrogen with the addition of 50% glycerol to prevent damage during long-term storage at −80°C.

For the preparation of full-length myosin from chemically skinned single *Musculus soleus* fiber, fibers were isolated from small bundles as described before (Kraft *et al*., 1995; Thedinga *et al*., 1999). Single fibers were sorted by their resting sarcomer length using laser light diffraction to ensure that only slow-twitch type-I fibers were used for myosin extraction. In brief, after isolation from bundles, muscle fibers were allowed to rest for 20 minutes at the bottom of a fiber preparation chamber under relaxing conditions at 5°C. Individual fibers were then placed in the centre of a 1.2 mW diode laser beam with a wavelength of 650 nm. The resulting diffraction pattern was projected onto a scaled analysis mask in a known distance below the preparation chamber bottom (Fig. S1). From the distance between main laser beam and the first maximum of diffracted laser light, the sarcomer length could be determined. Only fibers of resting sarcomer lengths of 2.0 μm or shorter were identified as slow-twitch type-I fibers and used for subsequent myosin extraction. Myosin from each fiber was extracted by placing the fiber attached to a glass capillary in a tube containing 5 μl extraction buffer for 30 min. Afterwards the fiber remains were removed by pulling the glass capillary out of the tube. The resulting myosin extracts were used immediately for *in vitro* motility assays. Remains of the single fiber myosin extracts were stored at −20°C for subsequent SDS-PAGE analysis of myosin light and heavy chain isoforms.

#### Unregulated actin filaments

Actin was prepared from chicken *pectoralis major* or rabbit back muscle as described previously (Pardee and Spudich, 1982), and fluorescently labelled, unregulated actin filaments were polymerized from tissue purified chicken pectoralis muscle G-actin as reported previously for rabbit back muscle actin (Scholz and Brenner, 2003). 0.12 nM chicken or rabbit G-actin were mixed with actin polymerisation buffer containing 10 mM HEPES pH 8, 100 mM KCl and 0.25 nM rhodamine phalloidin (Sigma-Aldrich, P1951). Unlabelled F-actin was polymerized accordingly, with the difference that 0.25 μM unlabelled phalloidin (Sigma-Aldrich, P2141) was used. Actin filaments were used for up to 3 days.

#### Regulated native thin filaments

Regulated thin filament were prepared from *soleus* or ventricular muscle tissue as described previously (Tobacman and Sawyer, 1990) and fluorescently labelled by the addition of 0.2 nM rhodamine-phalloidine. The thin filaments were stored on ice until use for up to 12 days.

### TIRF microscopy

#### Microscope image acquisition

Movement of fluorescently labelled actin or native thin filaments was recorded using a custom-made total internal reflection fluorescence (TIRF) microscope with single-fluorophore sensitivity (Amrute-Nayak *et al*., 2008; Rump *et al*., 2011) and modifications as described below. In our inverted objective-type TIRF microscope rhodamine phalloidin labelled actin or thin filaments were excited by light of 532 nm wavelength produced by a Nd:YAG laser (Compass 315M-150 SL, Coherent). Both exciting laser light and emitted fluorescence light passed through a high numerical aperture 60x oil immersion objective lens (Plan Apo 60x, NA 1.45, Oil; Olympus, Japan). Via a QV^2^ four-channel simultaneous-imaging system (Photometrics) with a respective dichroic mirror (T585 LPXR, Chroma) rhodamine fluorescence signals were projected onto a back-illuminated EMCCD camera (iXon DV887 Andor Technology, Belfast, UK, cooled to −80°C) and recorded with the software Andor SOLIS for imaging (Version 4.15.30000.0). The videos were recorded at a constant sample temperature of 23°C with a frame rate of 5 Hz and converted by Andor SOLIS to 16-bit grayscale tif-stacks for analysis with the computer program ImageJ (W.S. Rasband, National Institutes of Health, Bethesda, MD, USA) as described below.

#### *In vitro* motility

For all experiments with a 0.1% nitrocellulose-coated surface, first full-length myosin at a concentration of 1 mg/ml was immobilised on the surface for 2 min. After rinsing with motility extraction buffer (500 mM NaCl, 10 mM HEPES pH 7.0, 5 mM MgCl_2_, 2.5 mM MgATP), the surface was blocked with Pluronic F-127 (Sigma) for 1 min. Assay buffer AB (25 mM imidazole pH 7.4, 25 mM NaCl, 4 mM MgCl_2_, 1 mM EGTA, 10 mg/ml D-glucose, 10 mM DTT) was used to rinse the chamber before blocking of inactive myosin heads occurred for 1 min by unlabelled F-actin. AB containing additional 10 U/ml glucose oxidase, 800 U/ml catalase, and 2 mM ATP (A++) was flown into the chamber to release unlabelled actin from intact myosin heads. After a subsequent wash step with AB free of ATP (A+), fluorescently labelled F-actin was added to the chamber and incubated for 1 min. Unbound F-actin was rinsed out using A+. Filament gliding was initiated by adding A++ to the chamber and *in vitro* motility was immediately recorded. Temperature during recording was kept at 23°C. For experiments using regulated native thin filaments, A+, as well as A++ contained additional 1 mM CaCl_2_.

As an alternative to nitrocellulose 0.5 mg/ml BSA (Sigma-Aldrich, A6003) was used to coat the surface of the flow chamber for 5 min before full-length myosin at a concentration of 1 mg/ml was immobilised for 2 min. The chamber was rinsed with AB before inactive myosin was blocked using unlabelled F-actin for 1 min. A++ was used to release F-actin from active myosin heads. The chamber was once again washed with A+ before rhodamine phalloidin labelled F-actin or regulated native thin filaments were introduced for 1 min. Filament gliding and subsequent data recording was initiated using A++. Temperature during recording was kept at 23°C.

### Gel electrophoresis

#### Myosin heavy chain isoforms

For the separation of myosin heavy chain isoforms a large format SDS-PAGE setup with integrated cooling core was used (Protean II xi, Bio-Rad Laboratories, Inc.). A 5% stacking gel, containing 5% glycerol and a 6,5% separating gel containing 10% glycerol were cast using an acrylamide/bisacrylamide ratio of 30:0,3 (ROTIPHORESE^®^Gel 30, 3029.1; ROTIPHORESE^®^Gel A, 3037.1, Roth). The running buffer contained 0.0625 M Tris pH 8.3, 0.48 M glycine, 0.25% SDS and was precooled to 4 °C. Isoforms were separated at 10-25 mA for 32 h at 4 °C and subsequently stained using SYPRO® Ruby Protein Gel Stain (Supelco, S4942).

#### Myosin light chain isoforms

Myosin samples were separated using a small format 15% SDS-PAGE gel with a acrylamide/bisacrylamide ratio of 37.5:1 (ROTIPHORESE®Gel 30, 3029.1, Roth). Proteins were stained using Imperial™ Protein Stain (Thermo Scientific™), and subsequent densitometric analysis was carried out using an argus X1 gel documentation system (version 7.14.22, Biostep). Images were recorded as 16-bit grayscale tiff images, and proteins were quantified using Image Quant TL 1D (v.8.2.0.0, GE) software.

Western blot analyses of myosin light chain isoforms were performed after separation of proteins on a small format 12.5 % SDS-PAGE gel. For analysis of total protein, the nitrocellulose blotting membrane was stained with SYPRO®Ruby protein gel stain (Invitrogen™) prior to immunolabeling. Atrial and ventricular essential light chains MLC1a and MLC1s/v were immunolabelled using anti-MYL4 (Thermo Scientific™, PA5-49205). The atrial regulatory light chain (MLC2a) was labelled with a MLC2a specific antibody (Synaptic Systems GmbH, 311011).

#### Phospho-SDS-PAGE

For detection of phosphorylated light chains, extracted myosin samples were precipitated using ten times the volume of ultrapure water containing 2 mM DTT and 0.5 mM AEBSF.After incubation for 30 min on ice and centrifugation at 40000 rpm (Beckman Coulter, TLA 120.2), the pellet was resuspended in 1D sample buffer (62.5 mM Tris, pH 6.8, 15% Glycerol, 1% SDS, 0.002 % bromophenol blue) containing 1 tablet PhosSTOP™ (Roche, 4906837001) per 1ml. Proteins were separated using 12% Criterion™ TGX™ Precast Midi Protein Gel (Bio-Rad Laboratories, Inc., 5671043). The gel was first stained for phosphorylated proteins using Pro-Q™ Diamond Phosphoprotein Gel Stain (Molecular Probes® P33301) with PeppermintStick™ Phosphoprotein Molecular Weight Standards (Molecular Probes®, Inc., P27167) as a positive control. Subsequently, total protein was detected using SYPRO® Ruby Protein Gel Stain (Supelco, S4942). After densitometric analysis of the Pro-Q® Diamond signal (D) and the SYPRO® Ruby signal (S), the D/S ratios were calculated as a measure of phosphorylation (as described by manufacturer’s instructions; Molecular Probes®, Pro-Q™ Diamond Phosphoprotein Gel Stain).

### Data analysis and statistics

The velocity of fluorescently labelled actin or native thin filaments was manually analysed without any further image processing using the plug-in MTrackJ (Meijering *et al*., 2012) for the computer program ImageJ.

Velocity data was checked for normal distribution using Origin version 2020 (OriginLab Corporation, Northampton, MA, USA), which was also used to produce data plots. T-tests for significant differences between data sets were performed using the two sample, unequal variance TTEST function of Microsoft Excel 2016 (Microsoft Corporation, Redmond, WA, USA).

## Online supplemental material

Figure S1. **Determination of resting sarcomer length by laser diffraction.**

Figure S2. **SDS-PAGE analyses of actin and muscle thin filament preparations.**

Figure S3. **Sequence comparison of MLC1s isoforms.**

Figure S4. **Electrophoretic analysis of myosin light chain phosphorylation.**

Figure S5. ***M. soleus* single fiber actin gliding velocities (MLC1sa content of <40%, approx. 50%, and >60%, respectively).**

Table S1. **Mass spectrometry identification of MLC isoforms.**

Table S2. **Phosphorylation levels of *soleus* and ventricular myosin regulatory light chains.**

Movie S1. **Actin filament gliding on *soleus* (left) and ventricular (right) myosin preparations on a BSA coated chamber surface.**

## Results

### Gliding of actin filaments on ventricular myosin is significantly faster than on slow skeletal muscle myosin

Full-length ventricular β-/slow myosin molecules immobilized on a BSA coated assay chamber surface moved fluorescently labelled actin filaments with 0.865 ± 0.293 μm/s (Fig. 1A). This was significantly faster than actin filament gliding velocity on tissue purified full-length β-/slow *soleus* myosin (0.299 ± 0.131 μm/s, *p*=3.4*10^-205^) (Fig. 1A) (Supplemental movie S1).

**Figure 1.**
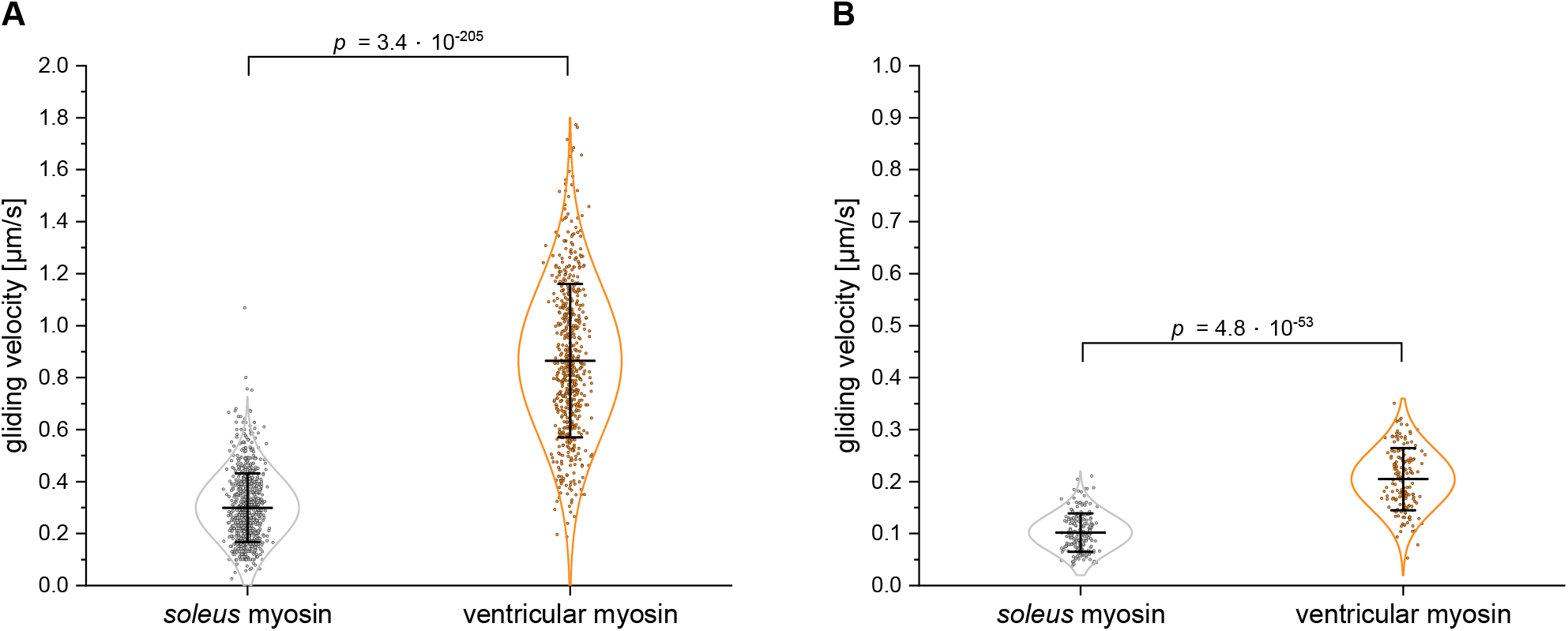
Gliding velocities of actin filaments on ventricular and *soleus* β-/slow muscle myosin. **(A)** On a BSA-coated assay chamber surface *soleus* β-/slow myosin molecules moved actin filaments with 0.299 μm/s (± 0.131 μm/s SD and 0.005 μm/s SE, n=821), while ventricular β-/slow moved actin filaments with 0.865 μm/s (± 0.293 μm/s SD and 0.012 μm/s SE, n=577). **(B)** On a nitrocellulose-coated assay chamber surface immobilized *soleus* β-/slow myosin molecules moved actin filaments with 0.102 μm/s (± 0.036 μm/s SD and 0.003 μm/s SE, n=196, 2 mM MgATP, T=23°C). Under the same conditions ventricular β-/slow moved actin filaments with 0.204 μm/s (± 0.059 μm/s SD and 0.005 μm/s SE, n=166).Data points represent gliding velocities of individual actin filaments, while bars represent mean values ± standard deviations. Data could be described by normal distributions (intrinsic curves).

The difference in gliding velocity was independent of the assay chamber coating and persisted on nitrocellulose-coated assay chamber surface (Fig. 1B). Overall, actin filament gliding on a nitrocellulose-coated surface was significantly slower compared to BSA-coated flow chambers (ventricular β-myosin *p*=2.3*10^-236^, *soleus* β-myosin *p*=9.3*10^-193^) (Fig. 1). Yet, ventricular myosin remained to move actin filaments significantly faster (0.204 ± 0.059 μm/s, *p*=4.8*10^-53^) than *soleus* myosin (0.102 ± 0.036 μm/s) (Fig.1B).

### Using native muscle thin filaments the differences in gliding speed between ventricular and *soleus* β-/slow myosin remains

Unregulated actin filaments are not the native interaction partner of striated muscle myosin as they lack regulatory proteins such as tropomyosin and the troponin complex. Consequently, the absence of actin-associated proteins might influence actomyosin interactions, and unregulated actin filaments are generally believed to slow down movement. However, compared to unregulated F-actin, tissue purified *soleus* thin filaments significantly decelerated movement on *soleus* myosin (0.082 ± 0.036 μm/s, *p*=3.9*10^-11^) (Fig. 2A). In contrast, thin filaments prepared from ventricular muscle moved significantly faster on ventricular β-myosin molecules (0.606 ± 0.254 μm/s, *p*=1.5*10^-215^) than unregulated actin filaments (Fig. 2B).

**Figure 2.**
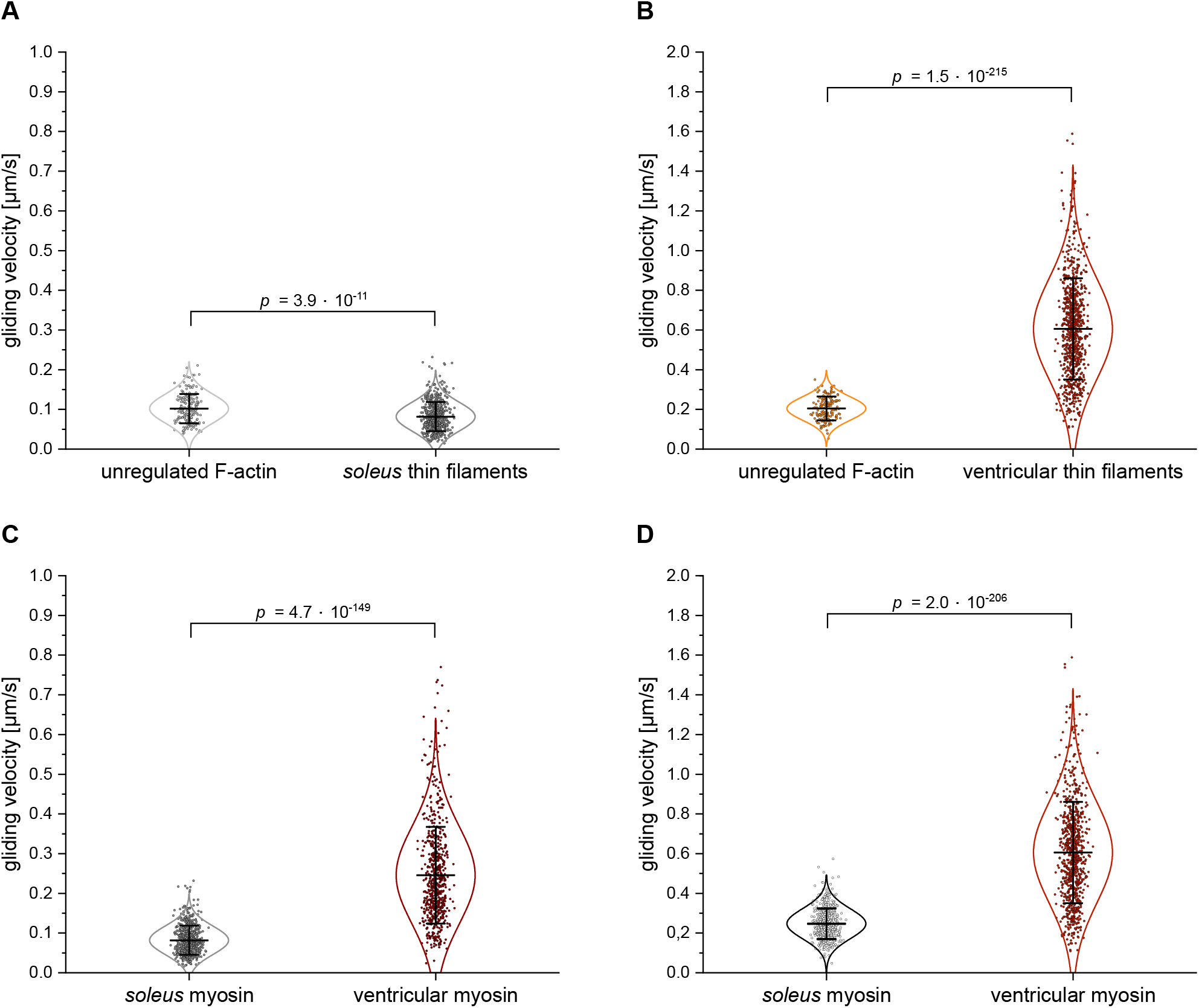
Gliding velocities of unregulated actin and native ventricular and *soleus* thin filaments on ventricular and *soleus* β-/slow muscle myosin on a nitrocellulose-coated assay chamber surface. **(A)** Movement of unregulated F-actin on *soleus* myosin (0.102 μm/s ± 0.036 μm/s SD and 0.003 μm/s SE, n=196) was significantly faster than of *soleus* native thin filaments (0.082 μm/s ± 0.036 μm/s SD and 0.001 μm/s SE, n=630). **(B)** Movement of unregulated F-actin on ventricular β-myosin molecules (0.204 μm/s ± 0.059 μm/s SD and 0.005 μm/s SE, n=166) was significantly slower than of thin filaments prepared from ventricular myofibrils (0.606 μm/s ± 0.254 μm/s SD and 0.009 μm/s SE, n=855). **(C)** Native thin filaments purified from *soleus* muscle were translocated significantly slower by *soleus* myosin (0.082 μm/s ± 0.036 μm/s SD and 0.001 μm/s SE, n=630, T=23°C) than by ventricular myosin (0.246 μm/s ± 0.121 μm/s SD and 0.005 μm/s SE, n=650). **(D)** *Soleus* myosin molecules moved cardiac thin filaments significantly slower (0.247 μm/s ± 0.076 μm/s SD and 0.003 μm/s SE, n=579) than ventricular myosin molecules (0.606 μm/s ± 0.254 μm/s SD and 0.009 μm/s SE, n=855). Data points represent gliding velocities of individual actin or thin filaments, while bars represent mean values ± standard deviations. Data could be described by normal distributions (intrinsic curves).

When using native thin filaments purified from *soleus* or ventricular myofibrils, the gliding velocities on *soleus* and ventricular β-/slow myosin on a nitrocellulose-coated surface remained significantly different. Movement of native thin filaments purified from *soleus* muscle on *soleus* was significantly slower (0.082 ± 0.036 μm/s, *p*=4.7 *10^-149^) than on ventricular myosin (0.246 ± 0.121 μm/s) (Fig. 2C). A similar effect was observed when cardiac native thin filaments were used. *Soleus* β-myosin translocated cardiac thin filaments with 0.247 ± 0.076 μm/s significantly slower compared to movement on ventricular β-myosin molecules (0.606 ± 0.254 μm/s, *p*=2.0*10^-206^) (Fig. 2D).

Differences in gliding speed between native thin filaments from ventricular or slow muscle tissue might arise from diverse muscle-type specific compositions of protein isoforms (Fig. S2). However, with all tested actin or native thin filament types significant differences in gliding velocity between *soleus* and ventricular β-/slow myosin molecules remained.

### Faster actin gliding on ventricular myosin cannot be explained by contaminations with faster atrial myosin

Both *soleus* and ventricular myosin contain the same myosin heavy chain (β-/slow-MyHC), encoded by gene *MYH7*. However, we found that myosin prepared from these two sources moved actin or native thin filaments with significantly different velocities, and the question remained whether *soleus* β-MyHC samples were decelerated compared to ventricular myosin or ventricular β-MyHC was accelerated by factors others than β-MyHC.

Since it has been reported that in some species ventricles might also express the faster atrial MyHC isoform α-MyHC in addition to β-MyHC to some extent (Sartore *et al*., 1981) or in response to increased levels of thyroid hormones (Lompré *et al*., 1984), we tested for possible contaminations with atrial myosin heavy or light chains in our myosin preparations. For rabbit ventricular myosin, no visible contamination with α-MyHC could be detected in myosin heavy chain isoform SDS-PAGE gels (Fig. 3A). While atrial myosin samples resolved in a double band representing the larger α-MyHC and slightly smaller β-MyHC isoform (Fig. 3A, lower panel), ventricular myosin showed only one band corresponding to β-MyHC (Fig. 3A, upper and lower panel). Contamination with atrial myosin light chains were also not detectable by SDS-PAGE (Fig. 3B) or by Western blot analysis (Fig. 3C). Neither antibodies raised against the atrial regulatory light chain MLC2a did detect any respective protein in ventricular myosin samples (Fig. 3C, lower panel), nor antibodies detecting both atrial and ventricular essential light chains MLC1a and MLC1v did suggest the presence of MLC1a in ventricular myosin samples (Fig. 3C, upper panel).

**Figure 3.**
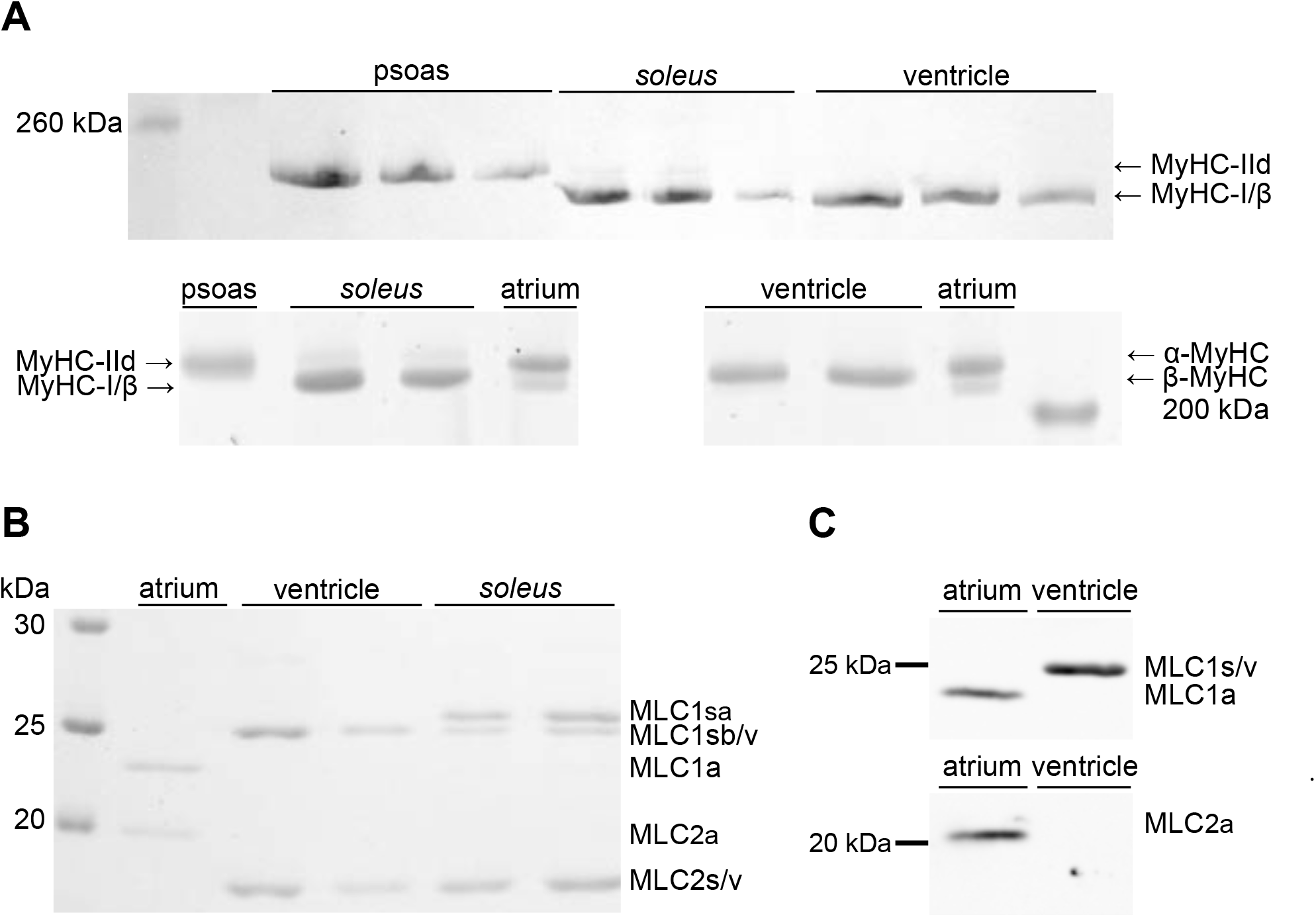
SDS-PAGE and Western blot analysis of ventricular and *soleus* muscle myosin heavy and light chain composition. **(A)** 8% (upper panel) and 6.5% (lower panels) SDS-PAGE analyses of myosin heavy chains in ventricular and *soleus* muscle myosin. Rabbit psoas muscle myosin containing MyHC-IId and atrial myosin containing both α- and β-MyHC served as references. No α-MyHC could be detected in myosin purified from ventricular tissue, while *soleus* muscle myosin contained traces of MyHC-IId (faint upper band). **(B)**15% SDS-PAGE analysis of myosin light chains in atrial, ventricular and *soleus* muscle myosin preparations. Atrial myosin contained the ELC MLC1a and RLC MLC2a, while ventricular myosin contained ELC MLC1sb/v and RLC MLC2s/v. In addition to the latter two light chains *soleus* muscle myosin contained the approx. 0.7 kDa larger ELC MLC1sa. **(C)** Western blot analysis of myosin light chains in atrial and ventricular myosin preparations. Both atrial and ventricular essential light chains MLC1a and MLC1s/v were immunolabelled by the antibody anti-MYL4 (upper panel), and no signal of MLC1a was detected in ventricular myosin preparations. The atrial regulatory light chain (MLC2a) was labelled with a MLC2a specific antibody (lower panel) and was absent in ventricular myosin samples.

**Figure 4.**
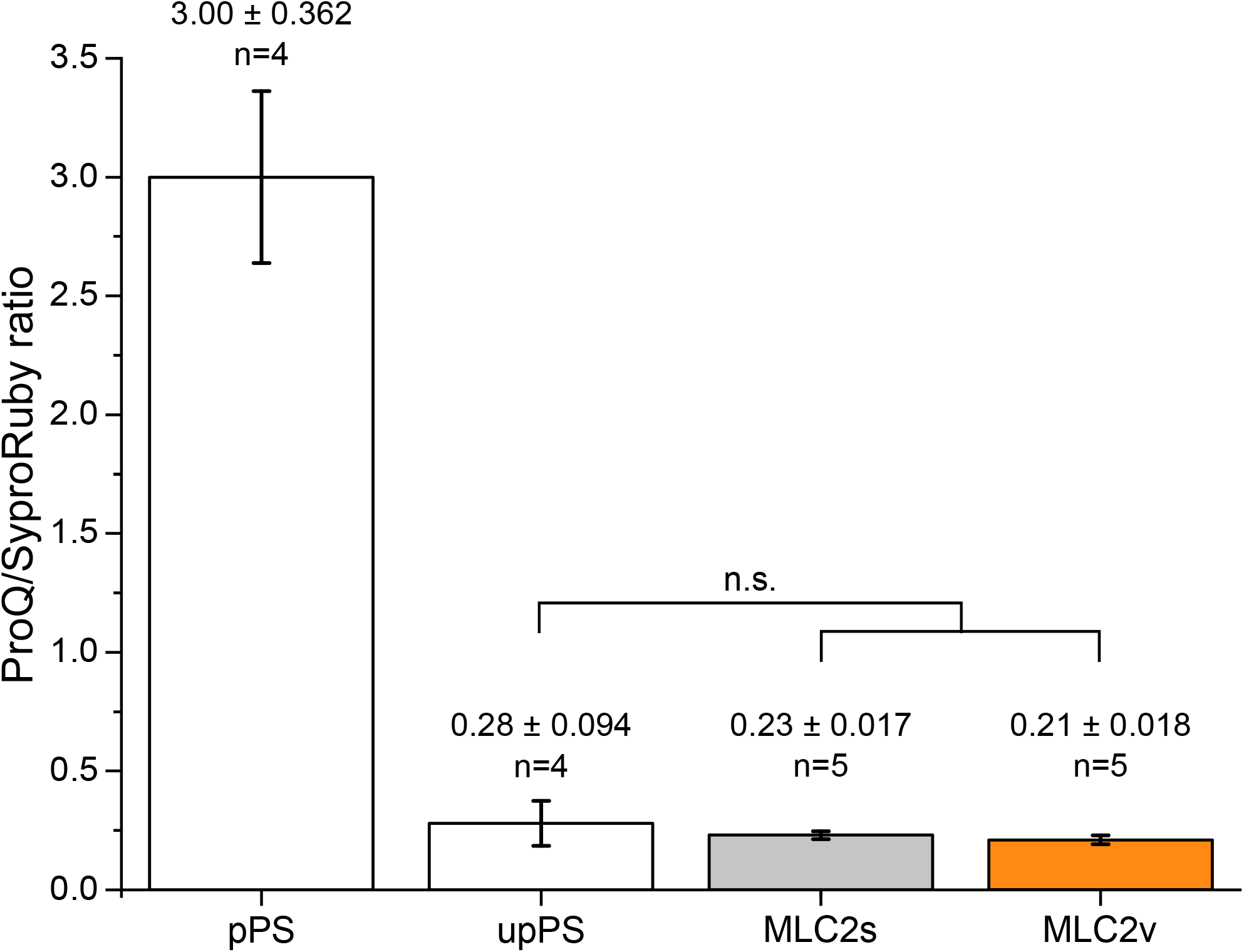
Phosphorylation levels of *soleus* and ventricular myosin regulatory light chains. Compared to a phosphorylation to protein fluorescence (Pro-Q/SyproRuby) ratio of the positive control (pPS, 45 kDa band of the PeppermintStick™ phosphoprotein standard) of 3.00 ±0.362 (SE, n=4), RLCs of *soleus* (MLC2s) and ventricular (MLC2v) myosin showed much lower and comparable ratios of 0.23 ±0.017 (SE, n=5) and 0.21 ±0.018 (SE, n=5), respectively. The unphosphorylated control protein (upPS, 116 kDa band of the PeppermintStick™ phosphoprotein standard) had a Pro-Q/SyproRuby fluorescence ratio of 0.28 ±0.094 (SE, n=4). Neither D/S ratios of *soleus* and ventricular RLCs were significantly different nor ratios of RLCs and of the unphosphorytated control protein. Bars represent mean values ± standard error.

### Although slow skeletal muscle myosin and ventricular myosin share the same myosin heavy chain they differ in their light chain composition

Immobilized *soleus* muscle β-/slow myosin molecules drove gliding of unregulated actin and native thin filaments with a lower velocity compared to ventricular β-/slow myosin, yet we could not detect any differences regarding their myosin heavy chains or contaminations with faster heavy or light chain isoforms in ventricular myosin preparations. However, from Fig. 3B it became obvious that they differed in their light chain compositions. *Musculus soleus* myosin contained, besides β-/slow-MyHC, slow/ventricular essential light chain MLC1sb/v and slow/ventricular regulatory light chain MLC2s/v, one additional, myosin associated protein migrating slightly slower than MLC2s/v. This protein was identified by mass spectrometry as the essential myosin light chain (ELC) isoform MLC1sa (Tab. S1), which was absent in ventricular myosin (Fig. 3B). Compared to MLC1sb/v, MLC1sa has a 13 amino acid larger N-terminal region (Fig. S3). This N-terminal region of long ELC isoforms has been reported to interact transiently with actin filaments (Milligan *et al*.,1990), thus serving as a second actin binding site of myosin molecules (Miyanishi *et al*.,2002; Nieznańska *et al*., 2002) and possibly modifying actomyosin interaction kinetics (Petzhold *et al*., 2014; Sweeney, 1995; VanBuren *et al*., 1994). Myosin preparations from *soleus* muscle tissue contained significant amounts of both ELC isoforms MLC1sa and MLC1sb with a higher relative content of MLC1sa (Fig. 3B).

Besides differences in myosin heavy or light chain isoform compositions, also post-translational modifications such as phosphorylation might affect actomyosin interactions thus leading to differences in actin gliding velocities. Standard SDS-PAGE and Western blot analysis do not exclude differences between *soleus* and ventricular myosin regarding the degree of phosphorylation e.g. of myosin regulatory light chains. Unlike in smooth muscle, regulatory light chain (RLC) phosphorylation does not act as an activation switch in striated muscle but rather modulates actomyosin interactions during striated muscle contraction. In skeletal muscle fibers, RLC phosphorylation accelerates actomyosin cross-bridge cycling kinetics by increasing the rate of non-force-generating to force-generating state transition (Sweeney and Stull, 1990). In contrast, *in vitro* motility assays showed that actin filament gliding slows down upon RLC phosphorylation due to an increase in the duty cycle of skeletal muscle myosin (Greenberg *et al*., 2009). Thus, to test for possible differences in the degree of RLC phosphorylation between our *soleus* and ventricular samples, which might influence actin-gliding velocity, we analysed *soleus* and ventricle muscle derived myosin samples accordingly. To perform an electrophoresis-based ratiometric analysis, we measured the intensity of the Pro-Q® Diamond phosphorylation signal (D) and the SYPRO® Ruby protein signal (S) of the respective protein bands, and calculated the D/S ratios of the fluorescence. We found that in *soleus* and ventricular myosin RLCs were, if at all, phosphorylated to very little and comparable degrees. Compared to a ratio of the positive control (45 kDa band of the PeppermintStick™ phosphoprotein standard) of 3.00 ± 0.362 (SE, n=4) phosphorylation to protein fluorescence, RLCs of *soleus* and ventricular myosin showed much lower but comparable ratios of 0.23 ± 0.017 (SE, n=5) and 0.21 ± 0.018 (SE, n=5), respectively (Tab. S2). The unphosphorylated control protein (116 kDa band of the PeppermintStick™ phosphoprotein standard) had a D/S fluorescence ratio of 0.28 ± 0.094 (SE, n=4). A picture of the respective gel is shown as supplemental figure S4.

### A higher content of the essential MLC1sa is associated with a reduced actin gliding velocity

Since myosin preparations from *soleus* muscle tissue contained significant amounts of both ELC isoforms MLC1sa and MLC1sb, we extracted myosin freshly from single *soleus* muscle fibers and tested for possible fiber-to-fiber variations regarding the MLC1sa/ MLC1sb ratio and respective actin gliding velocities on a BSA-coated assay chamber. All single fibers tested contained both ELC isoforms, most of them MLC1sa at a higher content (Fig. 5A). A higher relative content of MLC1sa compared to the “ventricular” ELC MLC1sb/v was associated with significantly slower actin gliding (grouping cut-off >60%: 0.311 ± 0.097 μm/s, and <40% MLC1sa: 0.393 ± 0.101 μm/s, respectively. *p*=3.3*10^-5^) (Fig. 5B), which suggests MLC1sa as the factor slowing down *soleus* compared to ventricular β-/slow-MyHC. In Fig. 5A, fibers 6 and 7 represent exemplary fibers containing >60% or <40% MLC1sa, respectively. Fibers containing MLC1sa and MLC1sb/v to equal amounts, exemplary represented by fiber 3 in Fig. 5A, translocated actin filaments at intermediate speed, not significantly different from fibers with a low relative content of MLC1sa, but significantly faster than fibers containing >60% MLC1sa (Fig. S5).

**Figure 5.**
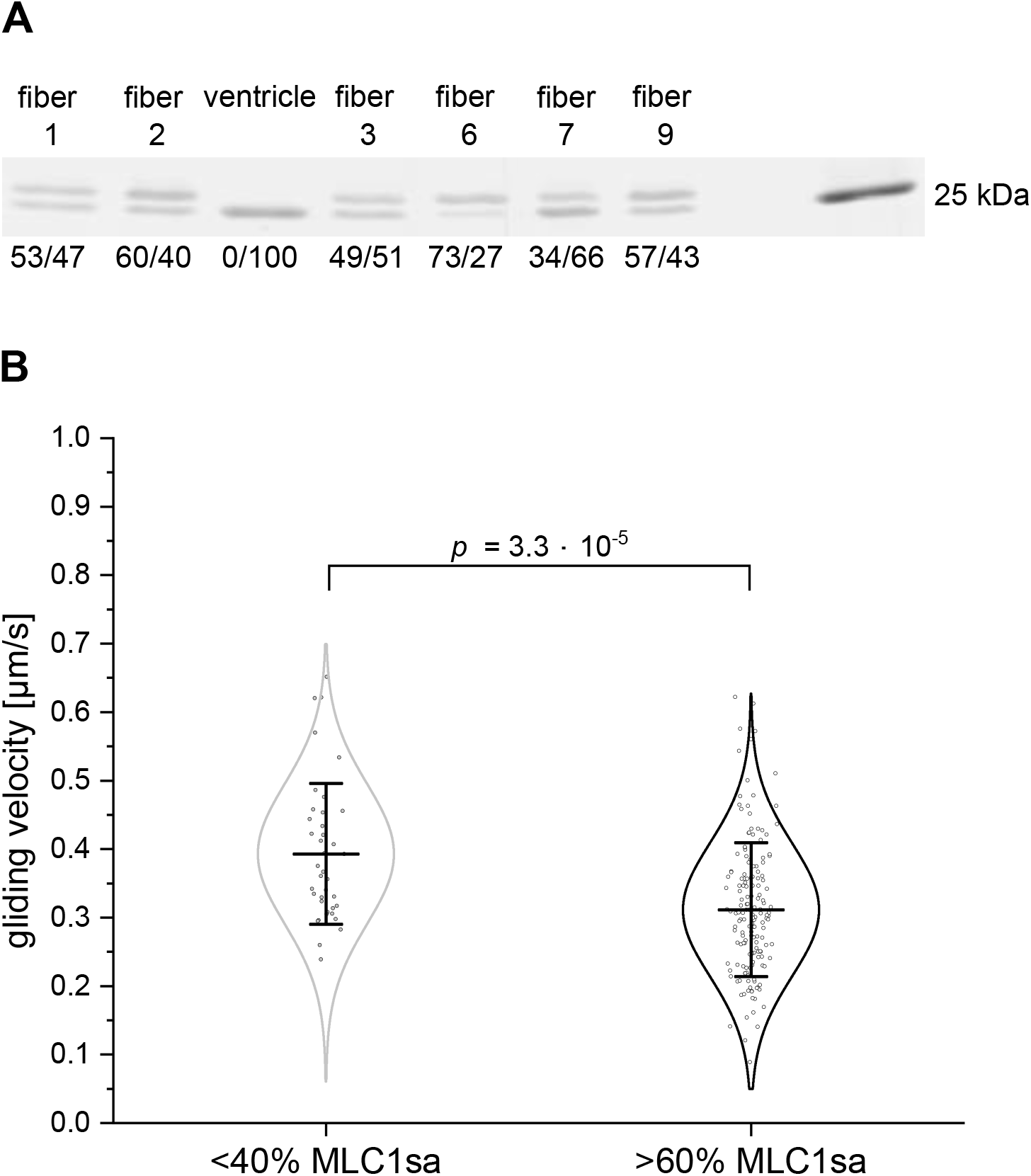
Myosin light chain composition and actin filament gliding velocity of individual type-I *soleus* muscle fibers. **(A)** Example of myosin light chain 15% SDS-PAGE analysis of individual type-I *soleus* muscle fibers. For each fiber the respective relative content of MLC1sa (upper band) versus MLC1sb/v (lower band) was determined by densitometric analysis. Note that all tested *soleus* fibers contained both ELC isoforms, the majority showed a higher content of MLC1sa, which was not detected in the control ventricular myosin. **(B)** On a BSA-coated assay chamber surface myosin extracted from individual type-I *soleus* fibers with a lower MLC1sa/ MLC1sb ratio (<40% MLC1sa) moved actin filaments with 0.393 μm/s (± 0.101 μm/s SD and 0.016 μm/s SE, n=39). Actin gliding on myosin prepared from *soleus* fibers containing >60% MLC1sa was significantly slower (0.311 μm/s ± 0.097 μm/s SD and 0.008 μm/s SE, n=166, T=23°C, *p*=3.3*10^-5^). Data points represent gliding velocities of individual actin filaments, while bars represent mean values ± standard deviations. Data could be described by normal distributions (intrinsic curves).

## Discussion

Many previous reports suggested that even when myosin molecules share the same motor heavy chain, their functional properties might differ due to differences in other components of the myosin complex such as associated light chains. In the present study we found that actin gliding velocity on ventricular myosin was significantly faster than on *M. soleus* myosin, although the MyHCs are identical. This difference was independent of assay chamber coating and persisted even when tissue purified native thin filaments were used. Faster actin gliding on ventricular myosin could not be explained by contaminations with faster atrial myosin heavy or light chains or by differences e.g. in RLC phosphorylation. However, slower actin gliding was associated with the presence of the additional essential light chain MLC1sa in *soleus* myosin, and on a single *soleus* muscle fiber level, a higher relative content of MLC1sa to MLC1sb/v was related with significantly slower actin gliding.

At saturating ATP concentrations, unloaded muscle shortening and actin gliding velocities, *V*, are thought to be limited by the detachment of actin-bound myosin heads that resist shortening (Huxley, 1957), and are therefore also related to the fraction of the time that a myosin head spends in its strongly-bound attached phase, the so-called duty ratio *r* (*r*=*τ_on_*/(*τ_on_*+ *τ_off_*)). Thus, according to this simple model actin gliding velocities are determined by *V*=d·*τ_on_^-1^* (Harada *et al*., 1990; Uyeda *et al*., 1990), where *d* is the displacement per actomyosin interaction (the myosin step size) and *τ_on_* is the lifetime of the attached states of the myosin heads. The rate of detachment of myosin from actin is limited by the rate of ADP release from actomyosin (Siemankowski and White, 1984; Warshaw *et al*., 1991; Yamashita *et al*., 1994) or the rate of attachment of ATP to actomyosin crossbridges (Nyitrai *et al*., 2006). However, besides speed limitations of unloaded muscle shortening by kinetics of myosin detachment from actin it also correlates with the maximal ATPase rate (Barany, 1967). Several studies also reported that unloaded velocity of fiber shortening exceeds the detachment limit (Baker *et al*., 2002; Haldeman *et al*., 2014; Hooft *et al*., 2007) and that factors that inhibit attachment kinetics slow down fiber shortening, too. Hence, it has been suggested that also attachment kinetics to actin might limit filament-sliding velocity at saturating myosin densities (Brizendine *et al*., 2015; Stewart *et al*., 2021). Consequently, possible explanations of a decreased actin gliding velocity in the present study might include (i) an increased duty ratio due to an increased actin affinity of myosin heads and lower detachment rates, (ii) slower attachment kinetics of myosin heads to actin, and/or (iii) a reduced myosin step size. Slower detachment kinetics would increase the time fraction of strongly-bound myosin states and thus the duty ratio of myosin, and might be the result of slower ATP binding to actomyosin or decelerated ADP release from actomyosin crossbridges.

In the present study, the presence of the additional essential myosin light chain MLC1sa in *soleus* myosin was associated with significantly slower actin gliding, and this deceleration was more pronounced at a higher relative content of MLC1sa. How could the presence of this, compared to ELC MLC1sb/v, longer and more positively charged ELC MLC1sa (Fig. S3) lead to one or more of the above mentioned decelerating causes? In early studies, it was shown that the unloaded shortening velocity of skinned skeletal muscle fibers as well as the velocity of actin filament gliding measured in *in vitro* motility assays depended on the ELC isoform composition of myosin. Fibers and extracted myosin with a high content of A1 type ELC LC1 shortened or moved actin filaments with a significantly lower velocity than those containing a high amount of short A2 type ELC LC3 (Greaser *et al*.,1988; Lowey *et al*., 1993a, b; Sweeney *et al*., 1988). Subsequently, many studies reported interactions of the “sticky” N-terminus of ventricular ELC (A1, *MYL3*) with actin. In an early cryoEM and traditional helical reconstruction study the decoration of actin filaments with myosin subfragment 1 (S1) either containing A1 type ELCs or and the shorter A2 type ELCs was compared. The resulting difference map showed an additional density peak at the C-terminal region of actin, which was assigned to the extra N-terminal residues in A1 (Milligan *et al*., 1990). This finding was consistent with a biochemical study showing that the N-terminal amino acids 1-15 of ventricular A1 ELCs could be cross-linked to acidic residues 361-364 of actin (Sutoh, 1982). It was also reported using myosin-induced actin polymerization experiments that myosin S1 containing N-terminal truncated A1 type ELCs lost the ability to interact with actin and to facilitate actin polymerization (Lowey *et al*.,2007). In striated muscle, A1 type ELCs can transiently interact with the negatively charged C-terminus of actin, resulting in strongly actin-bound myosin heads with slower kinetics compared to A2 ELCs (Hernandez *et al*., 2007; Morano, 1999; Sweeney, 1995). Weakening the A1/actin interaction by experimental interventions such as peptide and antibody-studies in skinned muscle fibers or N-terminal truncated A1 type ELCs *in vitro* increased myosin motor activity. The observed increased actin-activated myosin ATPase activity, faster *in vitro* actin motility speed, and higher maximal shortening velocity of skinned muscle fibers were interpreted as the result of an increased detachment rate of myosin from actin filaments (Bottinelli *et al*., 1994; Greaser *et al*., 1988; Hayashibara and Miyanishi, 1994; Lowey *et al*., 2007; Morano *et al*., 1995; Petzhold *et al*., 2014; Rarick *et al*., 1996; Sweeney, 1995; Timson *et al*., 1999; Wagner and Weeds, 1977).

The A1 ELC interaction with actin mainly resides in a cluster of conserved lysines K3, K4, K8, and K9 within its 41-43 residue N-terminal extension (Fodor *et al*., 1989; Seharaseyon *et al*., 1990) followed by a repeating sequence of proline/alanine residues. The overall 13 amino acid longer MLC1sa (*MYL6B*) contains K3, K4, and K9, too, plus two additional positive charges in very close vicinity at position 10 (Fig. S3). Compared to MLC1sb/v, MLC1sa is charged +2 within its first 13 N-terminal residues, which could enhance electrostatic interactions, thus, increasing the attachment rate and affinity to actin and decreasing the detachment rate from actin. Longer and more positively charged ELCs such as MLC1sa might, therefore, increase the duty ratio of myosin and therefore the load during filament sliding and lead to a lower velocity of actin filament gliding.

Besides interaction of long A1 type myosin ELCs with actin, also intramolecular interactions within the myosin motor domain have been reported. These intramolecular interactions involve the N-terminal extension of A1 ELC and the SH3 domain close to the nucleotide-binding site of the MyHC head domain, and it has been proposed that this might modulate ATPase kinetics, possibly slowing down ADP release or reducing the rate of ATP hydrolysis (Lowey *et al*., 2007). Hence, a longer and more positively charged N-terminal extension present in MLC1sa could slow down the release of ADP or decrease the rate of ATP hydrolysis even more, thus slowing down actin filament sliding. A transient A1/SH3 domain interaction could also bridge the gap between the ELC binding site at the lever arm of myosin and actin, thus, facilitating binding of the N-terminal extension to actin (Lowey *et al*., 2007).

In addition to actomyosin crossbridge kinetics, also the step size of myosin is one determinant of actin filament sliding velocity. Skeletal muscle myosin has a step size of approximately 5-6 nm (Steffen *et al*., 2001), while for cardiac myosin it has been suggested that an apparent step along actin might consist of 3 and 5 nm substeps of varying frequency, which eventually add up to a step size of 8 nm (Wang *et al*., 2016). It has also been proposed that the presence of the long N-terminal extension of ventricular A1 type myosin light chains can cause changes in the frequency of these subsequent substeps, and therefore of the step size. According to the proposed mechanism, binding of the N-terminal extension to the actin filament leads to an increased frequency of 5 nm steps followed by the additional 3 nm substep thus increasing step size. Yet, a larger step size alone would lead to faster actin sliding. Indeed, a higher frequency of total 8 nm steps was accompanied by lower ATPase rates and lower actin sliding velocities (Wang *et al*.,2016), which suggests a larger total step size was overcompensated by slower kinetics such as myosin detachment from actin. Compared to ventricular MLC1sb/v, the longer N-terminal extension of MLC1sa could lead to an even higher frequency of larger steps. Our observation that myosin containing higher amounts of MLC1sa moves actin and native thin filaments at decreased speeds, however, suggests that either step size is not increased or it is overcompensated by other decelerating processes such as slower crossbridge kinetics.

The additional myosin light chain MLC1sa found in *soleus* muscle fibers comprise an N-terminal extension, which is approximately 13 amino acids longer and more positively charged than MLC1sb/v found in ventricular myosin molecules. Following the idea that this might strengthen the interaction to actin and/or the SH3 domain of the myosin motor domain, what could be functional effects? Increased ELC/actin interactions could lead to (i) an altered step size, (ii) faster attachment kinetics, and (iii) an increased actin affinity with a reduced actin-activated ATPase and slower detachment kinetics. If the step size was changed, either it could be reduced to decelerate actin gliding to fit our observations, or, if increased following the model proposed by Wang and co-workers, this was overcompensated by a considerably reduced actin-activated ATPase presumably associated with a decreased actin detachment rate. Faster attachment kinetics of myosin to actin resulting from facilitated MLC1sa/actin interaction would lead to increased actin filament speeds, which does not fit our observations. Slower detachment kinetics could be the result of slower ATP binding to actomyosin, stronger and therefore prolonged binding of myosin heads to actin, or decelerated ADP release from actomyosin crossbridges. A decelerated release of ADP could be caused e.g. by intermolecular strain within the myosin ensemble or by intramolecular ELC/MyHC motor domain interactions. In experiments using myosin extracts from single *soleus* muscle fibers, a relative content of more than 60% of MLC1sa to MLC1sb/v was needed to decelerate actin filament sliding significantly. This suggests that in an ensemble of asynchronously working myosin molecules like in the *in vitro* motility assay, the presence of only few MLC1sa-containing myosin heads with decelerated actin detachment is not sufficient to slow down actin filaments. With increasing relative content of MLC1sa and therefore increased load on the ensemble crossbridges, load-dependent processes such as ADP release could be prolonged also in the primarily unaffected myosin heads, and actin-sliding velocity would be reduced.

We propose that the longer and more positively charged MLC1sa decelerates actin gliding either due to a higher affinity to actin associated with a decreased dissociation rate from actin compared to MLC1sb/v or by decelerated actomyosin cycling kinetics. Such ELC/actin interactions might also be relevant *in vivo* as differences between *soleus* and ventricular myosin remain when native thin filaments were used. To further test for possible MLC1sa-associated changes in attachment or detachment kinetics or in the step size of myosin single molecule studies such as optical trapping can be of great value in future studies.

## Acknowledgements

We are grateful to Bernhard Brenner (deceased) for helpful discussion in early stages of the study. The authors would like to thank the MHH Core Facility Proteomics for mass spectrometric analysis and identification of *soleus* myosin light chains. The authors also thank Judith Montag for alignment of light chain sequences and Torsten Beier for technical assistance. We thank Jan Faix for sharing muscle tissue. This work was supported by grants from the Young faculty program of Hannover Medical School (to Tim Scholz) and TW is supported by a research grant from Fritz Thyssen Stiftung (10.19.1.009MN to Mamta Amrute-Nayak). The authors declare no competing financial interests. All authors disclose any actual or potential conflict of interest including any financial, personal or other relationships with other people or organizations within three years of beginning the submitted work that could inappropriately influence, or be perceived to influence, their work.

## Author Contributions

Tim Scholz conceived the study, which was supervised by Tim Scholz, Mamta Amrute-Nayak, and Theresia Kraft. Jennifer Osten and Tim Scholz designed experiments. Maral Mohebbi, Jennifer Osten, Petra Uta, Tianbang Wang, and Tim Scholz performed experiments. Proteins were purified by Jennifer Osten, Faramarz Matinmehr, and Petra Uta. Jennifer Osten, Maral Mohebbi, Petra Uta, Mamta Amrute-Nayak, Tianbang Wang, and Tim Scholz analysed data. Jennifer Osten and Tim Scholz wrote the manuscript with contributions by Faramarz Matinmehr and revisions from Jennifer Osten, Mamta Amrute-Nayak, Theresia Kraft, and Tim Scholz.

## Abbreviations

TIRF: Total internal reflection fluorescence microscope
MyHC: microscope, myosin heavy chain
MLC: myosin light chain
ELC: essential light chain
RLC: regulatory light chain
HCM: hypertrophic cardiomyopathy

## References

Amrute-Nayak, M., Antognozzi, M., Scholz, T., Kojima, H., and Brenner, B. 2008. Inorganic Phosphate Binds to the Empty Nucleotide Binding Pocket of Conventional Myosin II*. Journal of Biological Chemistry 283, 3773–3781. https://doi.org/10.1074/jbc.M706779200

Amrute-Nayak, M., Nayak, A., Steffen, W., Tsiavaliaris, G., Scholz, T., and Brenner, B. 2019. Transformation of the Nonprocessive Fast Skeletal Myosin II into a Processive. Small, e1804313 LID - 1804310.1801002/smll.201804313 [doi].

Andreev, O.A., Saraswat, L.D., Lowey, S., Slaughter, C., and Borejdo, J. 1999. Interaction of the N-terminus of chicken skeletal essential light chain 1 with F-actin. Biochemistry 38, 2480–2485. 10.1021/bi981706x

Baker, J.E., Brosseau, C., Joel, P.B., and Warshaw, D.M. 2002. The biochemical kinetics underlying actin movement generated by one and many skeletal muscle Myosin molecules. Biophys J 82, 2134–2147.

Barany, M. 1967. ATPase activity of myosin correlated with speed of muscle shortening. J Gen Physiol 50,Suppl:197-218.

Barton, P.J., and Buckingham, M.E. 1985. The myosin alkali light chain proteins and their genes. Biochem J 231, 249–261. 10.1042/bj2310249

Becker, E., Navarro-López, F., Francino, A., Brenner, B., and Kraft, T. 2007. Quantification of mutant versus wild-type myosin in human muscle biopsies using nano-LC/ESI-MS. Analytical chemistry 79, 9531–9538. 10.1021/ac701711h

Bicer, S., and Reiser, P.J. 2004. Myosin light chain isoform expression among single mammalian skeletal muscle fibers: species variations. J Muscle Res Cell Motil 25, 623–633. 10.1007/s10974-004-5070-9

Biral, D., Ballarin, F., Toscano, I., Salviati, G., Yu, F., Larsson, L., and Betto, R. 1999. Gender- and thyroid hormone-related transitions of essential myosin light chain isoform expression in rat soleus muscle during ageing. Acta Physiol Scand 167, 317–323. 10.1046/j.1365-201x.1999.00621.x

Biral, D., Damiani, E., Volpe, P., Salviati, G., and Margreth, A. 1982. Polymorphism of myosin light chains. An electrophoretic and immunological study of rabbit skeletal-muscle myosins. Biochem J 203, 529–540. 10.1042/bj2030529

Bottinelli, R., Betto, R., Schiaffino, S., and Reggiani, C. 1994. Unloaded shortening velocity and myosin heavy chain and alkali light chain isoform composition in rat skeletal muscle fibres. J Physiol 478 (Pt 2), 341–349. 10.1113/jphysiol.1994.sp020254

Brizendine, R.K., Alcala, D.B., Carter, M.S., Haldeman, B.D., Facemyer, K.C., Baker, J.E., and Cremo, C.R. 2015. Velocities of unloaded muscle filaments are not limited by drag forces imposed by myosin crossbridges. Proc Natl Acad Sci U S A 112, 11235–11240. 10.1073/pnas.1510241112

Carraro, U., dalla Libera, L., and Catani, C. 1981. Myosin light chains of avian and mammalian slow muscles: evidence of intraspecific polymorphism. J Muscle Res Cell Motil 2, 335–342. 10.1007/bf00713271

Cuda, G., Fananapazir, L., Zhu, W.S., Sellers, J.R., and Epstein, N.D. 1993. Skeletal muscle expression and abnormal function of beta-myosin in hypertrophic cardiomyopathy. J Clin Invest 91, 2861–2865.

Cuda, G., Pate, E., Cooke, R., and Sellers, J.R. 1997. In vitro actin filament sliding velocities produced by mixtures of different types of myosin. Biophys J 72, 1767–1779.

Epstein, N.D., Cohn, G.M., Cyran, F., and Fananapazir, L. 1992. Differences in clinical expression of hypertrophic cardiomyopathy associated with two distinct mutations in the beta-myosin heavy chain gene. A 908Leu----Val mutation and a 403Arg----Gln mutation [see comments]. Circulation 86, 345–352.

Fodor, W.L., Darras, B., Seharaseyon, J., Falkenthal, S., Francke, U., and Vanin, E.F. 1989. Human ventricular/slow twitch myosin alkali light chain gene characterization, sequence, and chromosomal location. J Biol Chem 264, 2143–2149.

Geisterfer-Lowrance, A.A., Kass, S., Tanigawa, G., Vosberg, H.P., McKenna, W., Seidman, C.E., and Seidman, J.G. 1990. A molecular basis for familial hypertrophic cardiomyopathy: a beta cardiac myosin heavy chain gene missense mutation. Cell 62, 999–1006.

Greaser, M.L., Moss, R.L., and Reiser, P.J. 1988. Variations in contractile properties of rabbit single muscle fibres in relation to troponin T isoforms and myosin light chains. J Physiol 406, 85–98. 10.1113/jphysiol.1988.sp017370

Greenberg, M.J., Mealy, T.R., Watt, J.D., Jones, M., Szczesna-Cordary, D., and Moore, J.R. 2009. The molecular effects of skeletal muscle myosin regulatory light chain phosphorylation. American journal of physiology Regulatory, integrative and comparative physiology 297, R265–274. 10.1152/ajpregu.00171.2009

Guhathakurta, P., Prochniewicz, E., Roopnarine, O., Rohde, J.A., and Thomas, D.D. 2017. A Cardiomyopathy Mutation in the Myosin Essential Light Chain Alters Actomyosin Structure. Biophys J 113, 91–100. 10.1016/j.bpj.2017.05.027

Haldeman, B.D., Brizendine, R.K., Facemyer, K.C., Baker, J.E., and Cremo, C.R. 2014. The kinetics underlying the velocity of smooth muscle myosin filament sliding on actin filaments in vitro. J Biol Chem 289, 21055–21070. 10.1074/jbc.M114.564740

Harada, Y., Sakurada, K., Aoki, T., Thomas, D.D., and Yanagida, T. 1990. Mechanochemical coupling in actomyosin energy transduction studied by in vitro movement assay. J Mol Biol 216, 49–68.

Hayashibara, T., and Miyanishi, T. 1994. Binding of the amino-terminal region of myosin alkali 1 light chain to actin and its effect on actin-myosin interaction. Biochemistry 33, 12821–12827. 10.1021/bi00209a013

Hernandez, O.M., Jones, M., Guzman, G., and Szczesna-Cordary, D. 2007. Myosin essential light chain in health and disease. American journal of physiology Heart and circulatory physiology 292, H1643–1654. 10.1152/ajpheart.00931.2006

Hooft, A.M., Maki, E.J., Cox, K.K., and Baker, J.E. 2007. An accelerated state of myosin-based actin motility. Biochemistry 46, 3513–3520. 10.1021/bi0614840

Huang, W., and Szczesna-Cordary, D. 2015. Molecular mechanisms of cardiomyopathy phenotypes associated with myosin light chain mutations. J Muscle Res Cell Motil 36, 433–445. 10.1007/s10974-015-9423-3

Huxley, A.F. 1957. Muscle structure and theories of contraction. Prog Biophys Biophys Chem 7, 255–318.

Kirschner, S.E., Becker, E., Antognozzi, M., Kubis, H.P., Francino, A., Navarro-López, F., Bit-Avragim, N., Perrot, A., Mirrakhimov, M.M., Osterziel, K.J., et al. 2005. Hypertrophic cardiomyopathy-related beta-myosin mutations cause highly variable calcium sensitivity with functional imbalances among individual muscle cells. American journal of physiology Heart and circulatory physiology 288, H1242–1251. 10.1152/ajpheart.00686.2004

Köhler, J., Winkler, G., Schulte, I., Scholz, T., McKenna, W., Brenner, B., and Kraft, T. 2002. Mutation of the myosin converter domain alters cross-bridge elasticity. Proc Natl Acad Sci U S A 99, 3557–3562.

Kraft, T., Messerli, M., Rothen-Rutishauser, B., Perriard, J.C., Wallimann, T., and Brenner, B. 1995. Equilibration and exchange of fluorescently labeled molecules in skinned skeletal muscle fibers visualized by confocal microscopy. Biophys J 69, 1246–1258. 10.1016/S0006-3495(95)80018-4

Lompré, A.M., Nadal-Ginard, B., and Mahdavi, V. 1984. Expression of the cardiac ventricular alpha- and beta-myosin heavy chain genes is developmentally and hormonally regulated. J Biol Chem 259, 6437–6446.

Lowey, S., and Risby, D. 1971. Light chains from fast and slow muscle myosins. Nature 234, 81–85. 10.1038/234081a0

Lowey, S., Saraswat, L.D., Liu, H., Volkmann, N., and Hanein, D. 2007. Evidence for an interaction between the SH3 domain and the N-terminal extension of the essential light chain in class II myosins. J Mol Biol 371, 902–913. 10.1016/j.jmb.2007.05.080

Lowey, S., Waller, G.S., and Trybus, K.M. 1993a. Function of skeletal muscle myosin heavy and light chain isoforms by an in vitro motility assay. J Biol Chem 268, 20414–20418.

Lowey, S., Waller, G.S., and Trybus, K.M. 1993b. Skeletal muscle myosin light chains are essential for physiological speeds of shortening. Nature 365, 454–456.

Meijering, E., Dzyubachyk, O., and Smal, I. 2012. Methods for cell and particle tracking. Methods Enzymol 504, 183–200 LID - 110.1016/B1978-1010-1012-391857-391854.300009-391854 [doi].

Milligan, R.A., Whittaker, M., and Safer, D. 1990. Molecular structure of F-actin and location of surface binding sites. Nature 348, 217–221.

Miyanishi, T., Ishikawa, T., Hayashibara, T., Maita, T., and Wakabayashi, T. 2002. The two actin-binding regions on the myosin heads of cardiac muscle. Biochemistry 41, 5429–5438. 10.1021/bi0118355

Morano, I. 1999. Tuning the human heart molecular motors by myosin light chains. Journal of molecular medicine (Berlin, Germany) 77, 544–555. 10.1007/s001099900031

Morano, I., Ritter, O., Bonz, A., Timek, T., Vahl, C.F., and Michel, G. 1995. Myosin light chain-actin interaction regulates cardiac contractility. Circ Res 76, 720–725. 10.1161/01.res.76.5.720

Nier, V., Schultz, I., Brenner, B., Forssmann, W., and Raida, M. 1999. Variability in the ratio of mutant to wildtype myosin heavy chain present in the soleus muscle of patients with familial hypertrophic cardiomyopathy. A new approach for the quantification of mutant to wildtype protein. FEBS Lett 461, 246–252.

Nieznańska, H., Nieznański, K., and Stepkowski, D. 2002. The effects of the interaction of myosin essential light chain isoforms with actin in skeletal muscles. Acta Biochim Pol 49, 709–719.

Nyitrai, M., Rossi, R., Adamek, N., Pellegrino, M.A., Bottinelli, R., and Geeves, M.A. 2006. What limits the velocity of fast-skeletal muscle contraction in mammals? J Mol Biol 355, 432–442. 10.1016/j.jmb.2005.10.063

Pardee, J.D., and Spudich, J.A. 1982. Purification of muscle actin. Methods Enzymol 85 Pt B, 164–181.

Petzhold, D., Simsek, B., Meißner, R., Mahmoodzadeh, S., and Morano, I. 2014. Distinct interactions between actin and essential myosin light chain isoforms. Biochem Biophys Res Commun 449, 284–288. 10.1016/j.bbrc.2014.05.040

Pinter, K., Mabuchi, K., and Sreter, F.A. 1981. Isoenzymes of rabbit slow myosin. FEBS Lett 128, 336–338. 10.1016/0014-5793(81)80111-1

Rarick, H.M., Opgenorth, T.J., von Geldern, T.W., Wu-Wong, J.R., and Solaro, R.J. 1996. An essential myosin light chain peptide induces supramaximal stimulation of cardiac myofibrillar ATPase activity. J Biol Chem 271, 27039–27043. 10.1074/jbc.271.43.27039

Rayment, I., Holden, H.M., Sellers, J.R., Fananapazir, L., and Epstein, N.D. 1995. Structural interpretation of the mutations in the beta-cardiac myosin that have been implicated in familial hypertrophic cardiomyopathy. Proc Natl Acad Sci U S A 92, 3864–3868.

Rayment, I., Rypniewski, W.R., Schmidt-Base, K., Smith, R., Tomchick, D.R., Benning, M.M., Winkelmann, D.A., Wesenberg, G., and Holden, H.M. 1993. Three-dimensional structure of myosin subfragment-1: a molecular motor. Science 261, 50–58.

Reggiani, C., Potma, E.J., Bottinelli, R., Canepari, M., Pellegrino, M.A., and Stienen, G.J. 1997. Chemo-mechanical energy transduction in relation to myosin isoform composition in skeletal muscle fibres of the rat. J Physiol 502 (Pt 2), 449–460. 10.1111/j.1469-7793.1997.449bk.x

Reiser, P.J., and Bicer, S. 2006. Multiple isoforms of myosin light chain 1 in pig diaphragm slow fibers: correlation with maximal shortening velocity and force generation. Arch Biochem Biophys 456, 112–118. 10.1016/j.abb.2006.06.013

Reiser, P.J., Moss, R.L., Giulian, G.G., and Greaser, M.L. 1985. Shortening velocity in single fibers from adult rabbit soleus muscles is correlated with myosin heavy chain composition. J Biol Chem 260, 9077–9080.

Rump, A., Scholz, T., Thiel, C., Hartmann, F.K., Uta, P., Hinrichs, M.H., Taft, M.H., and Tsiavaliaris, G. 2011. Myosin-1C associates with microtubules and stabilizes the mitotic spindle during cell division. J Cell Sci 124, 2521–2528. Epub 2011 Jun 2528.

Sarkar, S., Sreter, F.A., and Gergely, J. 1971. Light chains of myosins from white, red, and cardiac muscles. Proc Natl Acad Sci U S A 68, 946–950. 10.1073/pnas.68.5.946

Sartore, S., Gorza, L., Pierobon Bormioli, S., Dalla Libera, L., and Schiaffino, S. 1981. Myosin types and fiber types in cardiac muscle. I. Ventricular myocardium. J Cell Biol 88, 226–233. 10.1083/jcb.88.1.226

Schiaffino, S., and Reggiani, C. 1996. Molecular diversity of myofibrillar proteins: gene regulation and functional significance. Physiol Rev 76, 371–423.

Scholz, T., and Brenner, B. 2003. Actin sliding on reconstituted myosin filaments containing only one myosin heavy chain isoform. J Muscle Res Cell Motil 24, 77–86.

Seebohm, B., Matinmehr, F., Köhler, J., Francino, A., Navarro-Lopéz, F., Perrot, A., Ozcelik, C., McKenna, W.J., Brenner, B., and Kraft, T. 2009. Cardiomyopathy mutations reveal variable region of myosin converter as major element of cross-bridge compliance. Biophys J 97, 806–824. 10.1016/j.bpj.2009.05.023

Seharaseyon, J., Bober, E., Hsieh, C.L., Fodor, W.L., Francke, U., Arnold, H.H., and Vanin, E.F. 1990. Human embryonic/atrial myosin alkali light chain gene: characterization, sequence, and chromosomal location. Genomics 7, 289–293. 10.1016/0888-7543(90)90554-8

Siemankowski, R.F., and White, H.D. 1984. Kinetics of the interaction between actin, ADP, and cardiac myosin-S1. Journal of Biological Chemistry 259, 5045–5053. https://doi.org/10.1016/S0021-9258(17)42953-X

Staron, R.S., and Pette, D. 1987. The multiplicity of combinations of myosin light chains and heavy chains in histochemically typed single fibres. Rabbit soleus muscle. Biochem J 243, 687–693. 10.1042/bj2430687

Steffen, W., Smith, D., Simmons, R., and Sleep, J. 2001. Mapping the actin filament with myosin. Proc Natl Acad Sci U S A 98, 14949–14954. Epub 12001 Dec 14944.

Stewart, T.J., Murthy, V., Dugan, S.P., and Baker, J.E. 2021. Velocity of myosin-based actin sliding depends on attachment and detachment kinetics and reaches a maximum when myosin-binding sites on actin saturate. J Biol Chem 297, 101178. 10.1016/j.jbc.2021.101178

Sutoh, K. 1982. Identification of myosin-binding sites on the actin sequence. Biochemistry 21, 3654–3661. 10.1021/bi00258a020

Sweeney, H.L. 1995. Function of the N terminus of the myosin essential light chain of vertebrate striated muscle. Biophys J 68, 112S–118S; discussion 118S-119S.

Sweeney, H.L., Kushmerick, M.J., Mabuchi, K., Sréter, F.A., and Gergely, J. 1988. Myosin alkali light chain and heavy chain variations correlate with altered shortening velocity of isolated skeletal muscle fibers. J Biol Chem 263, 9034–9039.

Sweeney, H.L., and Stull, J.T. 1990. Alteration of cross-bridge kinetics by myosin light chain phosphorylation in rabbit skeletal muscle: implications for regulation of actin-myosin interaction. Proceedings of the National Academy of Sciences of the United States of America 87, 414–418. 10.1073/pnas.87.1.414

Thedinga, E., Karim, N., Kraft, T., and Brenner, B. 1999. A single-fiber in vitro motility assay. In vitro sliding velocity of F-actin vs. unloaded shortening velocity in skinned muscle fibers. J Muscle Res Cell Motil 20, 785–796.

Timson, D.J., Trayer, H.R., Smith, K.J., and Trayer, I.P. 1999. Size and charge requirements for kinetic modulation and actin binding by alkali 1-type myosin essential light chains. J Biol Chem 274, 18271–18277. 10.1074/jbc.274.26.18271

Timson, D.J., Trayer, H.R., and Trayer, I.P. 1998. The N-terminus of A1-type myosin essential light chains binds actin and modulates myosin motor function. Eur J Biochem 255, 654–662. 10.1046/j.1432-1327.1998.2550654.x

Tobacman, L.S., and Sawyer, D. 1990. Calcium binds cooperatively to the regulatory sites of the cardiac thin filament. J Biol Chem 265, 931–939.

Tripathi, S., Schultz, I., Becker, E., Montag, J., Borchert, B., Francino, A., Navarro-Lopez, F., Perrot, A., Özcelik, C., Osterziel, K.J., et al. 2011. Unequal allelic expression of wild-type and mutated β-myosin in familial hypertrophic cardiomyopathy. Basic Res Cardiol 106, 1041–1055. 10.1007/s00395-011-0205-9

Uyeda, T.Q., Kron, S.J., and Spudich, J.A. 1990. Myosin step size. Estimation from slow sliding movement of actin over low densities of heavy meromyosin. J Mol Biol 214, 699–710. 10.1016/0022-2836(90)90287-v

VanBuren, P., Waller, G.S., Harris, D.E., Trybus, K.M., Warshaw, D.M., and Lowey, S. 1994. The essential light chain is required for full force production by skeletal muscle myosin. Proc Natl Acad Sci U S A 91, 12403–12407. 10.1073/pnas.91.26.12403

Wagner, P.D., and Weeds, A.G. 1977. Studies on the role of myosin alkali light chains. Recombination and hybridization of light chains and heavy chains in subfragment-1 preparations. J Mol Biol 109, 455–470. 10.1016/s0022-2836(77)80023-5

Wang, Y., Ajtai, K., Kazmierczak, K., Szczesna-Cordary, D., and Burghardt, T.P. 2016. N-Terminus of Cardiac Myosin Essential Light Chain Modulates Myosin Step-Size. Biochemistry 55, 186–198. 10.1021/acs.biochem.5b00817

Warshaw, D.M., Desrosiers, J.M., Work, S.S., and Trybus, K.M. 1991. Effects of MgATP, MgADP, and Pi on actin movement by smooth muscle myosin. J Biol Chem 266, 24339–24343.

Weeds, A.G. 1976. Light chains from slow-twitch muscle myosin. Eur J Biochem 66, 157–173. 10.1111/j.1432-1033.1976.tb10436.x

Yamashita, H., Sata, M., Sugiura, S., Momomura, S., Serizawa, T., and Iizuka, M. 1994. ADP inhibits the sliding velocity of fluorescent actin filaments on cardiac and skeletal myosins. Circ Res 74, 1027–1033.

Yu, Q.T., Ifegwu, J., Marian, A.J., Mares, A., Jr., Hill, R., Perryman, M.B., Bachinski, L.L., and Roberts, R. 1993. Hypertrophic cardiomyopathy mutation is expressed in messenger RNA of skeletal as well as cardiac muscle. Circulation 87, 406–412. 10.1161/01.cir.87.2.406

