## Supplemental material for "Myosin essential light chain 1sa decelerates actin and thin filament gliding on β-myosin molecules"

#### Author names and affiliations

Jennifer Osten[1,*], Maral Mohebbi[1,*], Petra Uta[1], Faramarz Matinmehr[1], Tianbang Wang[1], Theresia Kraft[1], Mamta Amrute-Nayak[1], and Tim Scholz[1,2]

[*] These co-first authors contributed equally to this work. [1]Molecular and Cellular Physiology, Hannover Medical School, Carl-Neuberg-Strasse 1, 30625 Hannover, Germany [2]To whom correspondence should be addressed: Tim Scholz, Tel: +49-511-5322737; Fax: +49-511-5324296;; Molecular and Cellular Physiology, Hannover Medical School, Carl-Neuberg-Strasse 1, 30625 Hannover, Germany, https://orcid.org/0000-0002-2127-9843.

#### Online supplemental material includes:

Figure S1. **Determination of resting sarcomer length by laser diffraction.**

Figure S2. **SDS-PAGE analyses of actin and muscle thin filament preparations.**

Figure S3. **Sequence comparison of MLC1s isoforms.**

Figure S4. **Electrophoretic analysis of myosin light chain phosphorylation.**

Figure S5. ***M. soleus* single fiber actin gliding velocities (MLC1sa content of <40%, approx. 50%, and >60%, respectively).**

Table S1. **Mass spectrometry identification of MLC isoforms.**

Table S2. **Phosphorylation levels of *soleus* and ventricular myosin regulatory light chains.**

Movie S1. **Actin filament gliding on *soleus* (left) and ventricular (right) myosin preparations on a BSA coated chamber surface.**

### Supplemental material

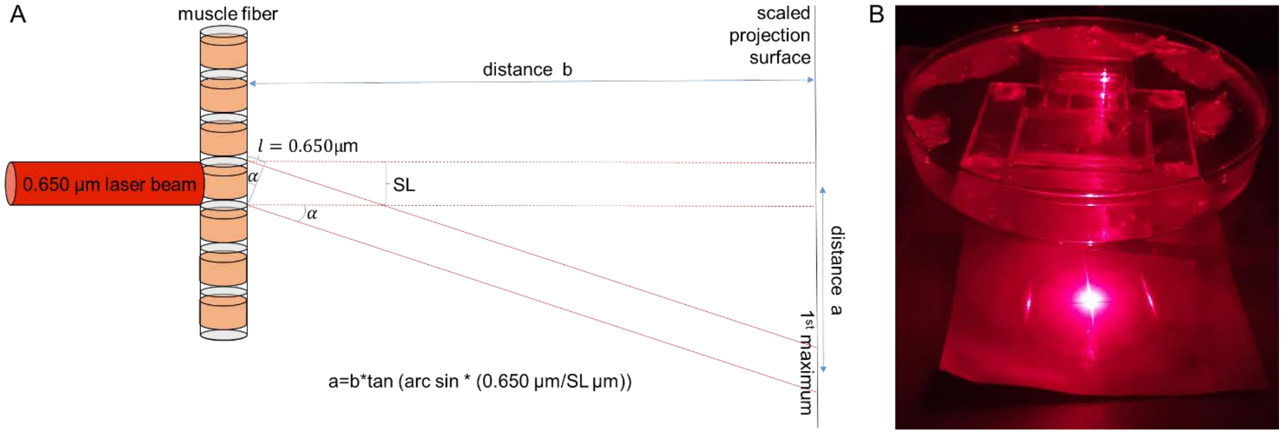


Figure S1. **Determination of resting sarcomer length by laser diffraction. (A)** Schematic drawing. **(B)** Photograph of the setup.

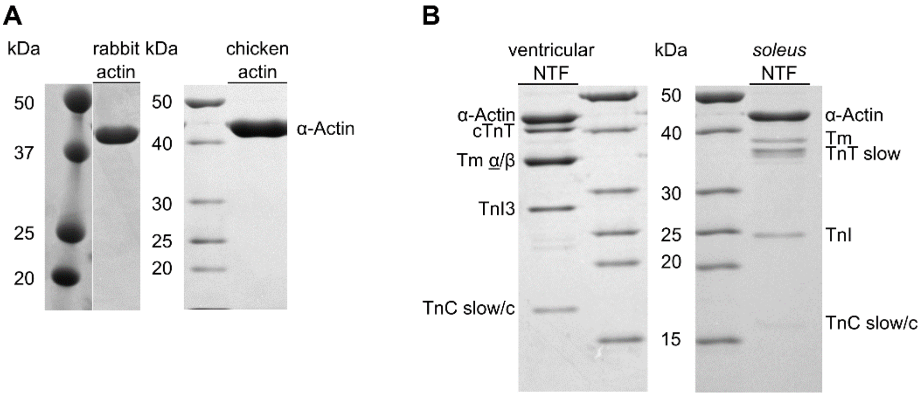


Figure S2. **SDS-PAGE analyses of actin and muscle thin filament preparations. (A)** Rabbit back muscle actin (Colormarker Biorad, #1610374, Precision Plus Protein Dual Color Standard) and chicken actin preparations (PageRuler™ Unstained Protein Ladder, ThermoScientific, #26614), and **(B)** ventricular (left) and *soleus* (right) muscle native thin filament composition.

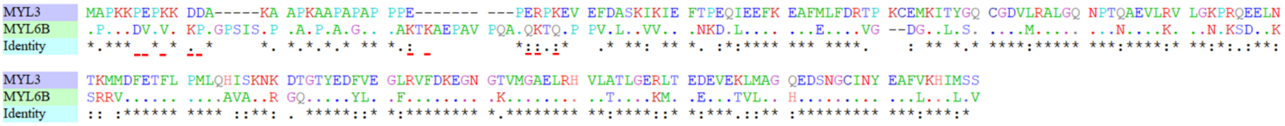


Figure S3. **Sequence comparison of MLC1s isoforms.** Accession No. for *MYL3* (MLC1sb/v) NP_000249 and for *MYL6B* (MLC1sa) NP_002466 were taken from NCBI. MYL3 (MLC1sb/v): 21932.05 Da, 195 AA; MYL6B (MLC1sa):  22763.98 Da, 208 AA. Red dashes indicate charge relevant changes and additional lysine residues within the N-terminal extensions.

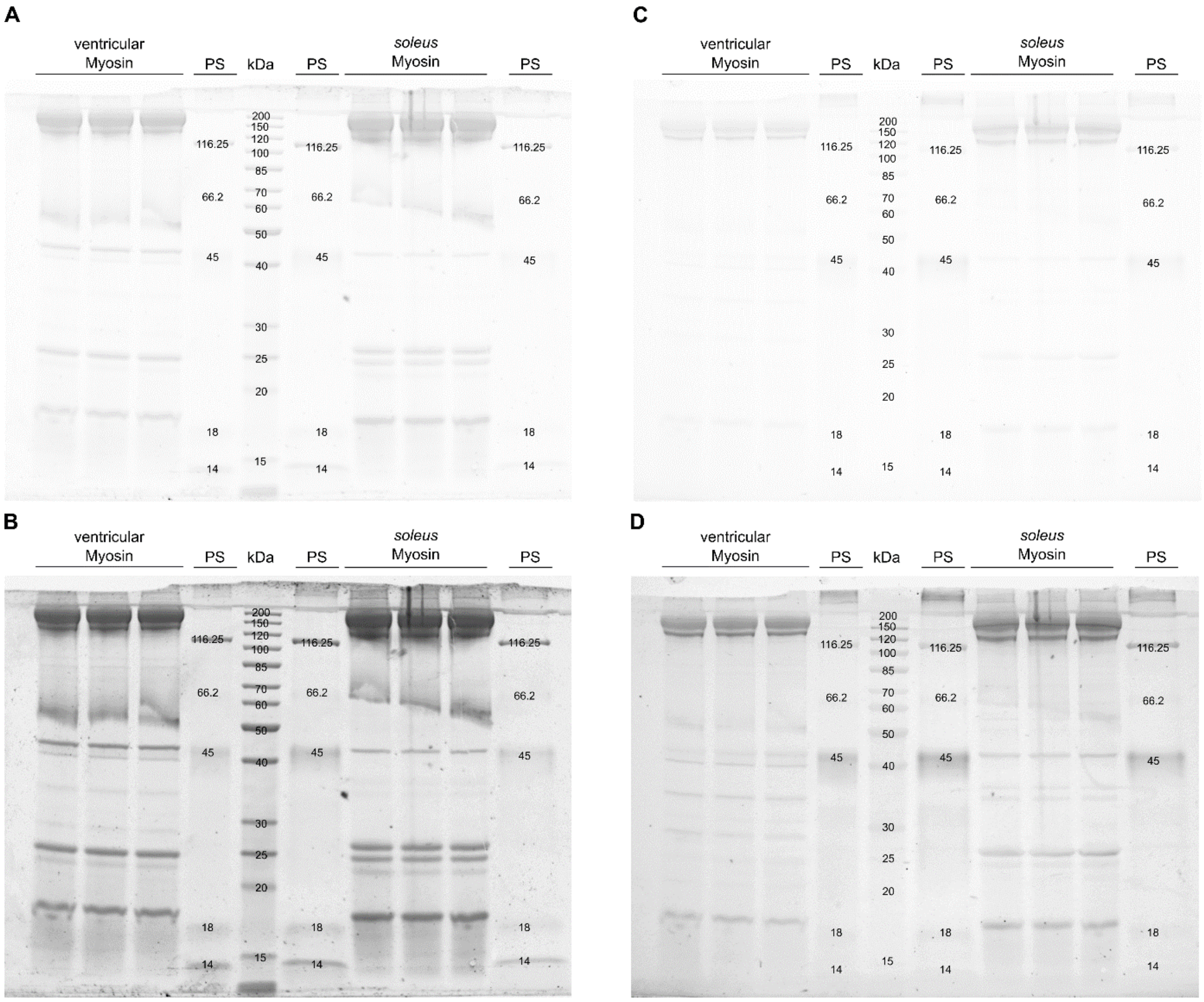


Figure S4. **Electrophoretic analysis of myosin light chain phosphorylation.** PS= PeppermintStick™ **(A)** SyproRuby, **(B)** SyproRuby colour corrected (-25 contrast, 0.22 gamma correction), **(C)** ProQ, and **(D)** ProQ colour corrected (-25 contrast, 0.22 gamma correction). Protein bands representing the regulatory light chains are found at a molecular weight of approximately 19 kDa.

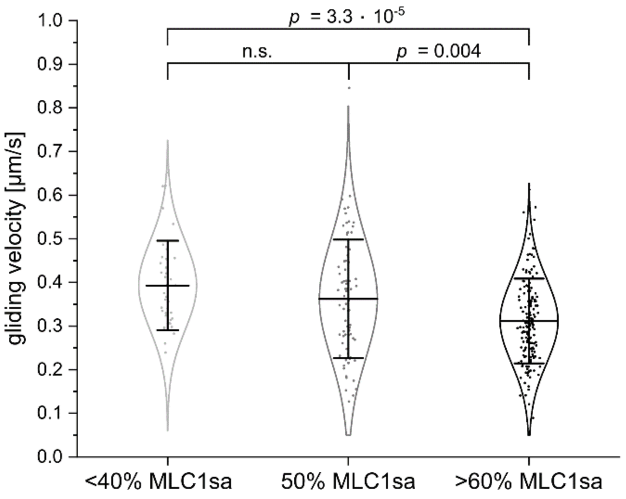


Figure S5. ***M. soleus* single fiber actin gliding velocities (MLC1sa contend of <40%, approx. 50%, and >60%, respectively).** On a BSA-coated assay chamber surface myosin extracted from individual type-I *soleus* fibers with a lower MLC1sa/MLC1sb ratio (<40% MLC1sa) moved actin filaments with 0.393 µm/s (± 0.101 µm/s SD and 0.016 µm/s SE, n=39). Actin gliding on myosin prepared from *soleus* fiber containing >60% MLC1sa was significantly slower (0.311 µm/s ± 0.097 µm/s SD and 0.008 µm/s SE, n=166, T=23°C, $p$=3.3*10$^{-5}$). Myosin extracted from individual type-I *soleus* fibers with equal amounts of MLC1sa and MLC1sb (50% MLC1sa) moved actin filaments with 0.362 µm/s (± 0.135 µm/s SD and 0.015 µm/s SE, n=76) significantly faster than myosin from fibers with a higher MLC1sa/MLC1sb ratio ($p$=0.004). Data points represent gliding velocities of individual actin filaments, while bars represent mean values ± standard deviations. Data could be described by normal distributions (intrinsic curves).

Table S1. **Mass spectrometric identification of MLC isoforms.**

| Accession | Description | Score | Coverage | # Proteins | # Unique Peptides | # Peptides | # PSMs | # AAs | MW [kDa] | calc. pI |
|---|---|---|---|---|---|---|---|---|---|---|
| A35-19_XP002713367 | myosin light chain 1sb-v Oryctolagus cuniculus | 2818,74 | 81,50 | 1 | 18 | 18 | 92 | 200 | 22,1 | 5,10 |
| FA35-18-NP002466.1 | myosin light chain 1sa Homo sapiens | 1143,12 | 60,58 | 1 | 17 | 17 | 58 | 208 | 22,7 | 5,73 |
| P13645 | Keratin | 863,22 | 32,04 | 28 | 13 | 16 | 30 | 593 | 59,5 | 5,21 |
| P35908 | Keratin | 860,63 | 39,22 | 5 | 17 | 20 | 28 | 645 | 65,8 | 8,00 |
| P04264 | Keratin | 704,88 | 32,30 | 11 | 17 | 20 | 27 | 644 | 66,0 | 8,12 |
| P00761 | Trypsin - Sus scrofa | 471,82 | 16,45 | 1 | 3 | 3 | 21 | 231 | 24,4 | 7,18 |
| P35527 | Keratin | 397,31 | 28,41 | 2 | 8 | 9 | 9 | 623 | 62,1 | 5,30 |
| P12883 | Myosin-7 Homo sapiens | 367,63 | 8,17 | 2 | 12 | 12 | 13 | 1935 | 223,0 | 5,80 |
| P02533 | Keratin | 334,33 | 13,35 | 34 | 3 | 6 | 11 | 472 | 51,6 | 5,16 |
| P13647 | Keratin | 229,48 | 12,20 | 7 | 5 | 7 | 8 | 590 | 62,3 | 7,74 |
| P02769 | Bovine serum albumin | 81,42 | 4,28 | 1 | 2 | 2 | 2 | 607 | 69,2 | 6,18 |
| FA84-17-ACTB-Variante-1 | ACTB Frameshift resulting in VLRVDRRLHPGLAVHLPADVDQQAGV | 66,68 | 4,49 | 3 | 1 | 1 | 1 | 356 | 39,4 | 5,47 |
| P02768-1 | Serum albumin | 62,19 | 4,93 | 1 | 2 | 2 | 2 | 609 | 69,3 | 6,28 |
| P62894 | Cytochrome c | 51,93 | 20,95 | 1 | 2 | 2 | 2 | 105 | 11,7 | 9,50 |
| Q5D862 | Filaggrin-2 | 39,83 | 0,50 | 1 | 1 | 1 | 1 | 2391 | 247,9 | 8,31 |
| Neat1-2 | FA42-17 ca 15kDa | 27,73 | 12,70 | 1 | 1 | 1 | 1 | 126 | 14,8 | 7,34 |
| O43790 | Keratin, type II | 20,74 | 1,44 | 3 | 1 | 1 | 1 | 486 | 53,5 | 5,66 |

Table S2. **Phosphorylation levels of *soleus* and ventricular myosin regulatory light chains.**

| **D/S ratio** | **45 kDa protein PeppermintStick™ (phosphorylation positive control)** | **116 kDa protein PeppermintStick™ (phosphorylation negative control)** | **M. *soleus* myosin regulatory light chain** | **ventricular myosin regulatory light chain** |
|---|---|---|---|---|
| **average** | 3.00 | 0.28 | 0.23 | 0.21 |
| **SD** | 0.723 | 0.189 | 0.038 | 0.039 |
| **SEM** | 0.362 | 0.094 | 0.017 | 0.018 |
| **n** | 4 | 4 | 5 | 5 |

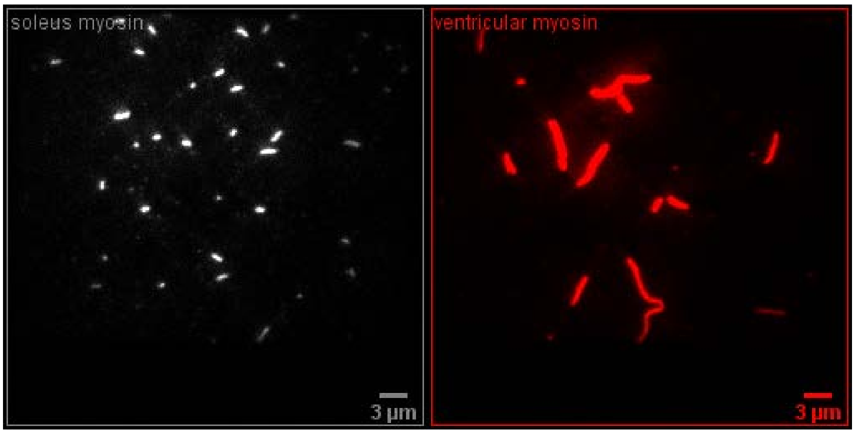

(first frame)

Movie S1. **Actin filament gliding on *soleus* (left) and ventricular (right) myosin preparations on a BSA coated chamber surface.** The average actin filament velocity on *soleus* myosin was 0.292 µm/s and 0.829 µm/s on ventricular myosin (T=23°C, 2 mM MgATP, played at speed of acquisition).
